# R-loop Prediction Reveals Generalization Limits of DNA Foundation Models Beyond Regulatory Genomics

**DOI:** 10.64898/2026.06.01.729367

**Authors:** Yafan Zhang, Arun Kumar Ganesan, Xingcheng Lin

## Abstract

DNA foundation models are increasingly proposed as general-purpose representations for genomic prediction and design, yet their evaluation remains largely centered on conventional regulatory tasks. This leaves a critical question unresolved: do DNA foundation models generalize to sequence biology beyond conventional gene regulation? To answer this question, we introduce **RloopBench**, a systematic benchmark for R-loop-forming sequence prediction as a biophysically distinct, genome-stability-associated task. We compare rule-based methods, task-specific models, classical sequence encodings, and foundation model representations across in-distribution, cross-platform, consensus-level, and cross-species evaluations. Foundation models achieve strong performance when positive and negative sequences are compositionally separable, but this advantage does not consistently transfer to cross-platform and cross-species settings, where they are often comparable to classical *k*-mer representations. Unexpectedly, a one-hot classifier baseline shows the strongest overall sensitivity to R-loop-forming sequences, exceeding more complex models across several generalization tests. Rule-based and task-specific models also exhibit limited transfer outside their original training regimes. Performance is further shaped by sequence properties, negative-control design, experimental platform, and species-specific genomic context. Together, RloopBench establishes genome-stability-associated sequence prediction as a complementary direction for DNA foundation model development and evaluation, while underscoring that simple sequence encodings remain necessary baselines for assessing model generalization beyond conventional regulatory tasks.

## 1 Introduction

Large-scale pretrained DNA foundation models have rapidly become important tools for genomic sequence analysis and design [1,2,3,4,5,6,7]. By learning from massive genomic corpora across diverse species, these models aim to encode reusable representations of sequence context, regulatory grammar, and long-range genomic dependencies. Recent DNA foundation models, including BERT-style masked language models [8,9], nucleotide transformer models [10], and long-context genomic sequence models [11,12], have been evaluated across a broad range of gene regulatory tasks [13,14]. These tasks include variant effect prediction, gene annotation tasks such as promoter and enhancer prediction, and epigenome-level annotations such as chromatin accessibility prediction and topologically associating domain (TAD) recognition. Although these benchmarks are biologically important, they represent only part of the functional genomic landscape.

An important open question is whether DNA foundation models can generalize beyond standard gene regulatory tasks. To address this question, we propose R-loop-forming sequence [15,16] prediction as a distinct genome stability-associated benchmark for evaluating whether DNA foundation models capture sequence features beyond conventional regulatory annotations. R-loops are three-stranded nucleic acid structures formed by an RNA–DNA hybrid and a displaced single-stranded DNA strand. Unlike standard regulatory annotations, R-loop formation is directly coupled to the biophysical properties [17,18,19,20] of nucleic acids, including RNA–DNA hybrid stability, strand asymmetry, G-richness, GC skew, and the potential formation of secondary structures such as G-quadruplexes. Thus, predicting R-loop-forming sequences requires models to capture sequence determinants related not only to regulatory grammar, but also to nucleic acid structure and genome stability. R-loops form co-transcriptionally and have been implicated in transcription regulation [16], replication stress [21], DNA damage [22], genome instability [23], and disease-associated genome dysfunction [24]. Therefore, R-loop prediction provides a biologically meaningful evaluation task for assessing whether pretrained DNA representations generalize from regulatory annotation tasks to biophysically grounded sequence features associated with genome stability.

Computational prediction of R-loop-forming sequences remains challenging. Previous approaches have used several different sequence representation strategies. Rule-based methods such as QmRLFS-finder [25] identify candidate R-loop-forming sequences using empirically derived sequence patterns, but may fail to detect R-loop sequences with noncanonical or previously uncharacterized features. Task-specific deep learning models, including DeepER [21] and deepRloopPre [26], use one-hot encoded DNA sequences as input to neural network architectures trained on experimentally mapped R-loop datasets. Although these models achieve good performance in their target datasets or species, their cross-platform and cross-species generalizability remains insufficiently evaluated. In addition, classical sequence representations such as *k*-mer frequencies have not been systematically benchmarked for R-loop prediction, despite their strong performance in many genomic sequence analysis tasks [27,28]. Given that DNA foundation models have shown promising cross-species generalization in other genomic applications, it is important to test whether similar behavior holds for R-loop prediction.

Here, we present RloopBench, a systematic benchmark of DNA sequence representations for R-loop prediction. We compare representative DNA foundation models, including Evo2, NTv3, and DNABERT-2, with classical sequence encodings, including 3-mer, 4-mer, and one-hot representations, as well as previously developed R-loop prediction methods such as QmRLFS-finder, DeepER, and deepRloopPre. The selected foundation models represent distinct architectural and pretraining designs, including StripedHyena-based long-context genomic modeling, U-Net-like nucleotide representation learning, and BERT-style masked language modeling. To ensure that performance differences primarily reflect the quality of the sequence representations rather than downstream classifier complexity, we evaluated newly considered representations using a unified linear-probe [29] framework.

We evaluate these methods across multiple complementary benchmarks, including the original DeepER R-ChIP dataset [21], an independent DRIPc-seq validation dataset [25,30], R-loopBase consensus zones [31] supported by multiple experimental technologies, and curated cross-species R-loop datasets. This design enables us to assess both cross-platform and cross-species generalizability. We find that DNA foundation model embeddings perform extremely well on the DeepER R-ChIP benchmark, where positive and negative sequences show significant sequence-level differences. However, this advantage does not consistently transfer to independent, cross-platform, or cross-species settings. Unexpectedly, the one-hot baseline achieves the best balanced classification performance on the independent DRIPc-seq dataset and maintains the highest recall across R-loopBase consensus zones and cross-species datasets. Classical *k*-mer methods are generally competitive with the long-context Evo2 representation and show stronger generalizability than NTv3 and DNABERT-2 in several evaluations. We further quantify sequence-level properties across datasets to interpret these benchmark patterns.

Our analyses show that R-loop prediction performance is strongly influenced by sequence composition, negative-control design, experimental platform, and species-specific genomic context. These results highlight the importance of careful dataset construction, especially the definition of true non-R-loop-forming sequences, for future R-loop predictor development. They also show that DNA foundation models should be evaluated beyond standard regulatory tasks, and that simple, biologically interpretable sequence encodings remain competitive for genome stability-associated sequence prediction.

This study makes four main contributions: **(A)** introducing R-loop prediction as a genome stability-associated benchmark beyond conventional gene regulatory tasks; **(B)** systematically comparing rule-based methods, classical encodings, task-specific deep learning models, and pretrained foundation model embeddings; **(C)** evaluating model behavior across in-distribution, independent cross-platform, multi-level consensus, and cross-species R-loop benchmarks; and **(D)** showing that DNA foundation models perform strongly on datasets with clear sequence-level separation but do not consistently outperform simple sequence encodings under more stringent validation settings.

## 2 Materials and Methods

### 2.1 DNA Sequence Representation

As summarized in Table 1, we considered multiple DNA sequence representation strategies for R-loop prediction. Their underlying principles and implementation details are described below.

**Table 1:** Representative paradigms for DNA sequence representation in computational R-loop prediction.

| Paradigm | Representative Models | Underlying Logic |
| --- | --- | --- |
| I. Rule-based pattern matching | QmRLFS-finder | Detects sequence motifs consistent with canonical R-loop-forming sequences, including a G-cluster-rich *R-loop initiation zone* (RIZ), a structurally unspecified linker region, and a downstream G-dense *R-loop elongation zone* (REZ). |
| II. Classical sequence encoding | $k$-mer (3/4-mer); one-hot + linear probe (baseline) | Represents DNA sequences using local compositional features (e.g., $k$-mer frequency vectors) or position-wise one-hot encoding, followed by a simple linear classifier for discrimination. |
| III. Task-specific deep learning | DeepER; deepRloopPre | Learns hierarchical spatial features directly from one-hot encoded DNA sequences using convolutional neural networks (CNNs) and recurrent architectures such as (bi)LSTMs. |
| IV. Genomic foundation models | Evo2; DNABERT-2; NTV3 embeddings + linear probe | Utilizes large-scale pretrained DNA foundation models to generate contextual sequence embeddings that capture long-range genomic dependencies, followed by a lightweight linear classifier. |

#### 2.1.1 Rule-based pattern matching

We evaluated QmRLFS-finder [25], a rule-based method based on the RIZ–Linker–REZ structural model for identifying potential R-loop-forming sequences (RLFS). The model detects G-rich sequence patterns associated with R-loop initiation and elongation regions. We applied the generalized model (Rules m1 and m2) to both DNA strands using default parameters to identify RLFS within each input sequence (see http://r-loop.org/?pg=qmrlfs). Since QmRLFS-finder reports RLFS at the subsequence level, a sequence was considered a positive prediction if at least one RLFS was detected within the sequence.

#### 2.1.2 Classical sequence encoding

We considered classical sequence encoding methods, including *k*-mer-based representations (3-mer and 4-mer) and one-hot encoding, as illustrated in Fig. 1A and B, respectively. The resulting feature vectors were subsequently used as input to the linear probe classifier (Section 2.2.1).

**Fig. 1:**
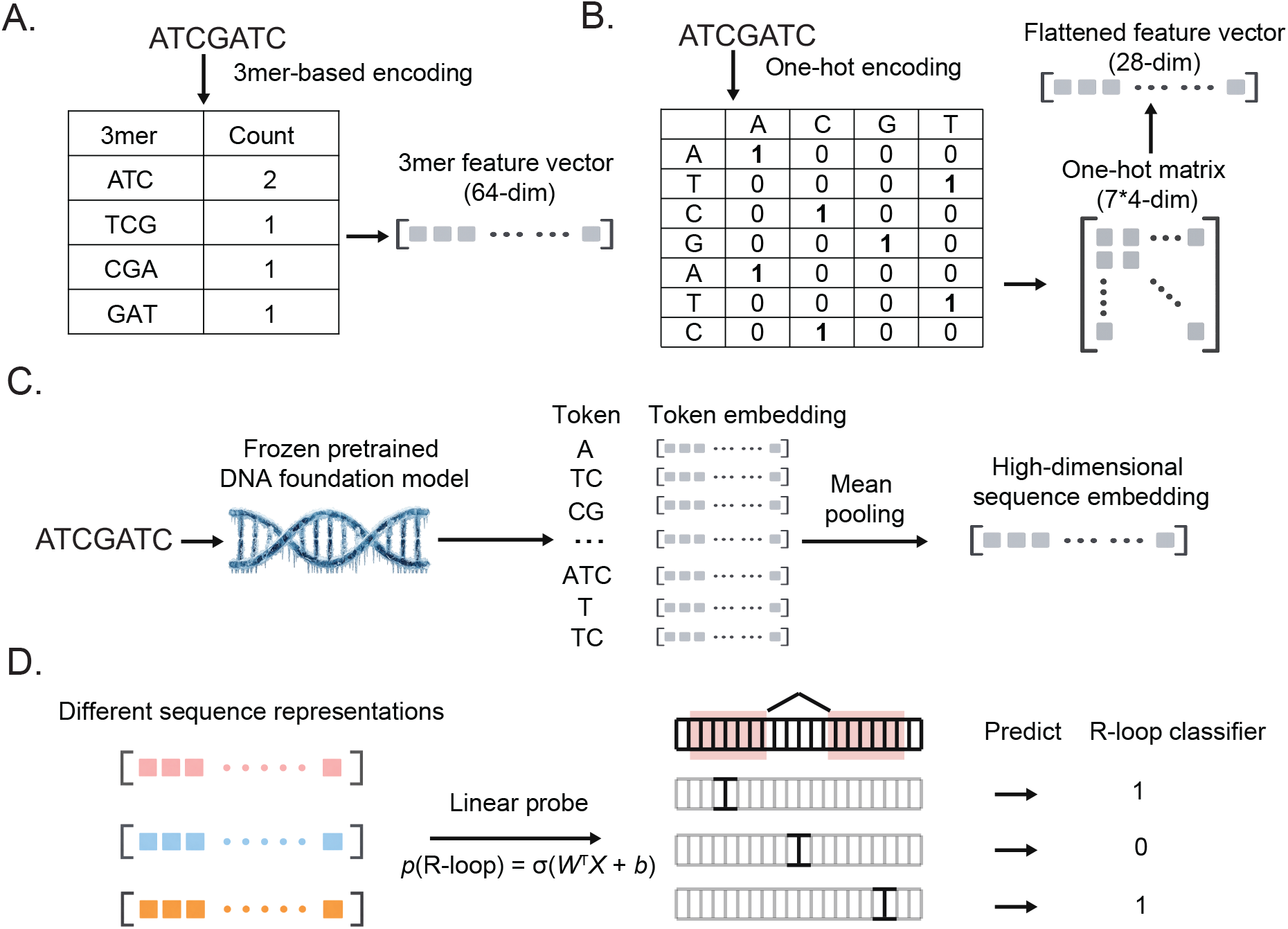
Schematic overview of DNA sequence representation strategies and the R-loop prediction task studied in this work. **(A)** *k***-mer-based representation:** Each DNA sequence is decomposed into overlapping *k*-mers and represented as a frequency-based feature vector. For example, 3-mer and 4-mer representations yield 64-dimensional and 256-dimensional feature vectors, respectively. **(B) One-hot representation:** Each nucleotide is encoded as a binary vector, preserving the positional composition of the input sequence. These vectors are then flattened into a single feature vector for downstream classifier development. **(C) DNA foundation model representation:** Input sequences are processed by a frozen pretrained DNA foundation model. The definition of a token differs across models: Evo2 represents sequences at single-nucleotide resolution, NTv3 uses near-single-nucleotide-resolution tokens with model-specific padding requirements, whereas DNABERT-2 uses BPE tokens that can correspond to variable-length DNA subsequences. Hidden states from a selected layer are used as token-level representations and aggregated by mean pooling to generate a sequence-level embedding, with dimensionality depending on the foundation model. **(D) Downstream classification task:** Given an input DNA sequence, the classifier outputs the probability of R-loop formation.

For the *k*-mer representation, each DNA sequence was encoded as a normalized *k*-mer frequency vector. Specifically, all possible combinations of A, C, G, and T were enumerated, resulting in a 64-dimensional feature vector for *k* = 3 and a 256-dimensional feature vector for *k* = 4. *k*-mers were extracted from each sequence using a sliding window, counted, and converted into relative frequencies by normalizing with the total number of valid observed *k*-mers. *k*-mers containing ambiguous bases (e.g., N) were excluded. This representation is independent of sequence length.

For one-hot encoding, sequence length was standardized because the representation is position-wise and therefore produces matrices whose dimensions depend directly on input sequence length. To ensure that all sequences could be represented with a fixed input dimension for downstream linear-probe evaluation, each sequence was adjusted to 5,000 bp by center-cropping sequences longer than 5,000 bp and symmetrically padding shorter sequences with N characters so that the original sequence remained centered. The processed sequences were then transformed into one-hot matrices, where A, C, G, and T were encoded as four-dimensional binary vectors, and ambiguous bases such as N were encoded as all-zero vectors.

#### 2.1.3 Task-specific deep learning

We evaluated two related published methods, DeepER [21] and deepRloopPre [26]. Both models take one-hot encoded DNA sequences as input and aim to capture sequence-dependent features associated with R-loop formation. They share a common design principle of combining convolutional layers for local pattern extraction with bidirectional LSTM (BiLSTM) modules to model long-range dependencies along the sequence.

However, they differ substantially in training data and prediction strategy. DeepER was trained on human R-ChIP-mapped R-loop peaks. Based on its codebase (https://github.com/NjuChenlab/DeepER), we first used `script.py` to generate base-level probability profiles. Following the authors’ original pipeline, we applied `def_rloop.py` to obtain region-level predictions using a sliding-window aggregation scheme with a window size of 200 bp and a step size of 10 bp. Regions with an average probability greater than 0.95 were considered positive predictions, following the optimal cutoff reported in the original DeepER study.

In contrast, deepRloopPre provides four pretrained models based on ssDRIP-seq data from *Arabidopsis*, zebrafish, human, and rice (http://bioinfor.kib.ac.cn/R-loopAtlas/l2/deep_downloads.html). For evaluation, we used the corresponding species-specific model when available; otherwise, the human model was applied. The deepRloopPredict.py script (https://github.com/PEHGP/deepRloopPre) directly outputs probabilities for non-overlapping 128 bp windows, and windows with an average probability greater than 0.5 (default cutoff) were considered positive predictions.

#### 2.1.4 Genomic foundation models

To comprehensively evaluate DNA foundation models, we selected three recent genomic language models that represent distinct state-of-the-art design choices: Evo2 [11], Nucleotide Transformer version 3 (NTv3) [2], and DNABERT-2 [9]. These models take DNA sequences as input, tokenize them into discrete units, and generate high-dimensional contextual embeddings for each token through multiple transformation layers (Fig. 1C).

To ensure consistent evaluation across models, all DNA foundation model embeddings were processed using a unified inference and prediction framework. For each model, token-level embeddings were first summarized into fixed-length window-level representations using model-specific pooling strategies. For sequences shorter than or equal to the model-specific window size, a single embedding was generated directly from the full sequence and passed to the linear probe classifier (Section 2.2.1). For longer sequences, we applied a sliding-window strategy: each window was independently processed by the corresponding foundation model, converted into a fixed-length embedding, and passed through the same linear probe classifier to obtain a prediction probability. The final sequence-level score was computed by averaging prediction probabilities across all windows. The model architectures and customized embedding extraction procedures are described below.

Evo2 is trained on 9 trillion DNA base pairs from a highly curated genomic database, OpenGenome2, spanning all domains of life. It is built upon the StripedHyena 2 architecture [32], a convolutional multi-hybrid autoregressive model specifically designed to handle long-context sequences (up to 1 million tokens) while capturing single-nucleotide resolution signals. Based on the official implementation (https://github.com/arcinstitute/evo2), we used the evo2_7B model, which provides a balance between predictive performance and computational efficiency. This version is numerically stable, supporting inference in bfloat16 precision on standard GPUs without requiring a Transformer Engine. We extracted embeddings from the recommended layer (blocks.28.mlp.l3). For each input window, token-level representations were generated with a dimensionality of 4,096 per token. A sliding window of 5,000 bp with a step size of 500 bp was applied across each sequence. The resulting token-level embeddings were mean-pooled across the sequence dimension to obtain a single 4,096-dimensional vector for each window. NTv3 was pretrained on the same datasets as Evo2 but adopts a distinct U-Net-like [33] masked modeling architecture to achieve comparable tokenization resolution and long-context modeling capability. We used the largest available NTv3_650M_pre model via the official Hugging Face implementation (https://huggingface.co/InstaDeepAI/NTv3_650M_pre). Because NTv3 requires input sequence lengths to be divisible by 128 due to its underlying Flash Attention mechanism, sequences were processed using a sliding window approach with a window size of 5,120 bp and a step size of 512 bp. Each window was padded to the nearest multiple of 128 prior to tokenization. For each processed window, we extracted the final hidden states from the last transformer layer of NTv3. Each valid token was represented by a 1,536-dimensional vector, corresponding to the model’s hidden size. These token-level embeddings were then summarized via masked mean pooling over valid token positions, using the attention mask to exclude dynamically padded positions and produce a fixed-length representation for each window.

DNABERT-2 is a BERT-style [34] transformer model that replaces the traditional *k*-mer tokenization used in DNABERT [8] with a Byte Pair Encoding (BPE) scheme, enabling more flexible and data-driven representation of DNA sequences. It is trained using a masked language modeling objective to capture contextual dependencies. The model was pretrained on both the human reference genome (2.75 billion bp) and a large multi-species genomic dataset comprising 135 species across diverse taxonomic groups, totaling 32.49 billion bp. We used the DNABERT-2-117M model via the official implementation (https://github.com/MAGICS-LAB/DNABERT_2). Sequences were processed using a sliding window of 5,000 bp with a step size of 500 bp. For each window, token-level embeddings (768 dimensions) were extracted from the final hidden layer. To obtain a fixed-length representation, embeddings were mean-pooled across valid tokens using the attention mask, thereby excluding padded positions from the pooled sequence embedding.

### 2.2 Training, Evaluation, and Testing

#### 2.2.1 Linear Probe Classifier

To isolate the contribution of different sequence representations, we employed a linear probe classifier [29] for methods newly considered in this study (Section 2.1.2 and 2.1.4), while previously published methods (Section 2.1.1 and 2.1.3) were evaluated using their original implementations. Specifically, given an input vector **x** *∈* ℝ^*d*^, the model computes a scalar logit:

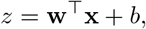

where **w** ∈ ℝ ^*d*^ and *b* ∈ ℝ are learnable parameters. The prediction probability is then obtained via the sigmoid function:

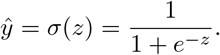

The input dimensionality *d* depends on the representation method, including classical feature vectors (e.g., 64 for 3-mer, 256 for 4-mer, and 20,000 for one-hot representations) and foundation model-derived embeddings (e.g., 768 for DNABERT-2, 1,536 for NTv3, and 4,096 for Evo2). Accordingly, the total number of trainable parameters in the linear probe is *d* + 1, reflecting the weight vector and bias term.

This simple architecture ensures that performance differences primarily reflect the quality of sequence representations rather than additional gains from further multi-layer optimization.

#### 2.2.2 Training Setup

To ensure a fair comparison, all models developed in this study were trained and evaluated on the same data and with the same training/validation/testing split as the original DeepER framework [21], using a 7:2:1 ratio. Positive and negative samples have the same number in each split. Input representation vectors were standardized using z-score normalization, with the scaler fitted on the training set and then applied to the validation and test sets.

The linear probe classifier was trained using the Adam optimizer with binary cross-entropy loss with logits. Let *z* denote the model output (logit), and *ω*(*z*) the sigmoid function. The loss is defined as:

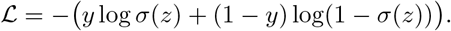

During inference, probabilities *ŷ* = *ω*(*z*) were thresholded at 0.5 to obtain binary predictions. Hyperparameters were selected by grid search over learning rates {10^*−*4^, 10^*−*3^, 10^*−*2^}and weight decay values {0, 10^*−*4^, 10^*−*2^ }. For each hyperparameter combination, the model was trained for up to 100 epochs, and validation performance was monitored after each epoch. Early stopping with a patience of 10 epochs was applied based on the validation F1-score, which was used because it jointly accounts for precision and recall and is therefore suitable for evaluating binary R-loop classification performance. The model achieving the highest validation F1-score across all hyperparameter configurations was selected for final evaluation. The F1-score is defined as:

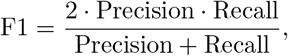

where precision and recall are given by:

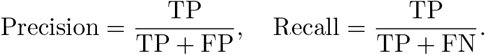

Here, TP, TN, FP, and FN represent true positives, true negatives, false positives, and false negatives, respectively.

#### 2.2.3 Evaluation Metrics

For the datasets described in Sections S1.1.1 and S1.1.2, which contain both positive sequences, defined as sequences capable of forming R-loops, and negative sequences, defined as sequences not observed to form R-loops, model performance was evaluated using standard classification metrics, including Accuracy, Precision, Recall, F1-score, Specificity, ROC-AUC, and Precision–Recall AUC (PR-AUC). Accuracy and Specificity were calculated as follows:

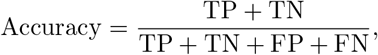

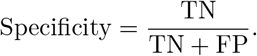

ROC-AUC was used to measure the overall ability of each model to distinguish positive from negative sequences across decision thresholds, whereas PR-AUC was used to evaluate the precision–recall tradeoff, particularly under class-imbalanced settings.

For the remaining datasets (Sections S1.1.3 and S1.1.4), where comparable negative sequences are difficult to define, we considered only recall to evaluate whether each model could capture R-loop-forming sequences.

### 2.3 Sequence Property Calculation

To characterize sequence-level properties across R-loop datasets, we computed composition, strand-asymmetry, G-richness, and sequence-complexity features for each subset. The resulting dataset-level summaries are reported in Table 4. All analyses were performed on processed sequences containing only standard nucleotides. For each sequence in a subset, let *L* denote the sequence length, and let *A, C, G*, and *T* denote the corresponding nucleotide counts.

GC content and G content were calculated as

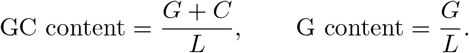

Strand asymmetry was quantified using GC skew and AT skew:

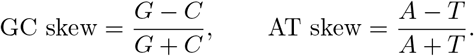

CpG density was defined as the frequency of the dinucleotide “CG”:

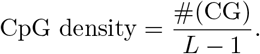

G-run density was used to quantify clustered G-rich sequence patterns and was defined as the number of continuous *G*_*≥*3_ runs per kilobase:

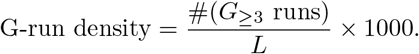

Following the canonical putative G-quadruplex sequence (PQS) definition used for genome-wide G4 motif prediction [35], potential G-quadruplex-forming sequences were identified using the pattern

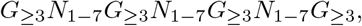

with an additional constraint that each loop segment contained only non-G nucleotides. The G4 motif fraction was calculated as the fraction of sequences in each subset containing at least one such motif.

Sequence complexity was measured using normalized Shannon entropy:

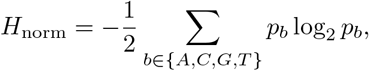

where *p*_*b*_ is the frequency of nucleotide *b*. The normalization factor of 2 corresponds to the maximum entropy for four equally frequent nucleotides, so *H*_norm_ ranges from 0 to 1.

For each subset, sequence-level values were aggregated by the mean unless otherwise specified. GC content, G content, CpG density, and G4 motif fraction were reported as percentages; GC skew, AT skew, G-run density, and normalized Shannon entropy were reported directly.

## 3 Results

### 3.1 Benchmark design for comparing DNA sequence representations in R-loop prediction

Through a literature review, we identified three representative methods for R-loop prediction that were previously developed: QmRLFS-finder [25] (with a web-based implementation known as R-loop tracker [30]), deepRloopPre [26], and DeepER [21]. QmRLFS-finder is a rule-based method based on empirically derived R-loop-forming sequence patterns, whereas deepRloopPre and DeepER use one-hot encoded DNA sequences as input to task-specific neural network models. The architectural and usage details of these methods are described in Sections 2.1.1 and 2.1.3.

In addition to these existing R-loop prediction methods, we included several newly evaluated representation strategies. First, because previous R-loop prediction benchmarks have not systematically considered traditional k-mer representations, we evaluated 3-mer and 4-mer frequency vectors as classical sequence-composition baselines (Fig. 1A; Section 2.1.2). Second, we added a one-hot baseline to test how much predictive signal can be recovered from position-wise sequence information without using a deep neural network architecture (Fig. 1B; Section 2.1.2). Third, we evaluated embeddings from pretrained DNA foundation models (Fig. 1C; Section 2.1.4) from Evo2 [11], NTv3 [2], and DNABERT-2 [9]. These models were selected because they are widely recognized representative DNA foundation models and differ in architecture, sequence context length, and pretraining strategy. Together, they cover complementary design choices, including StripedHyena-based [32] long-context genomic modeling in Evo2, U-Net-like [33] nucleotide representation learning in NTv3, and BERT-style [34] masked language modeling in DNABERT-2. This selection allowed us to test whether different forms of large-scale genomic pretraining provide additional predictive power for R-loop-associated sequence features (Section 2.1.4).

Based on these considerations, we organized all evaluated methods into four representative paradigms of DNA sequence representation: rule-based pattern matching, classical sequence encodings, task-specific deep learning models, and pretrained DNA foundation model embeddings (Table 1). This organization allowed us to compare methods that differ not only in model complexity, but also in the biological assumptions encoded in their input representations. Rule-based methods explicitly encode canonical R-loop sequence grammar. Classical encodings mainly capture local nucleotide composition and position-wise sequence patterns. Task-specific deep learning models learn discriminative features from labeled R-loop datasets. Foundation models, in contrast, rely on large-scale pretraining to capture broader genomic sequence dependencies.

We treated R-loop prediction as a binary classification task, in which each DNA sequence was classified as either R-loop-forming or non-R-loop-forming (Fig. 1D). For newly evaluated representations, we used a linear probe classifier [29] to minimize the contribution of downstream model complexity (Section 2.2.1). This design was intended to ensure that benchmark performance primarily reflects the quality of the input sequence representation rather than extensive deep classifier optimization.

For initial supervised training, validation, and testing, we used the R-ChIP dataset originally adopted in the DeepER study. We chose this dataset for two main reasons. First, it provides a direct compari-son with DeepER, which was previously reported to outperform R-loop Tracker, QmRLFS-finder, and deepRloopPre on this benchmark. Second, the dataset was derived from strand-specific R-ChIP-mapped R-loops in HEK293 and K562 cells and was further filtered using additional R-loop mapping evidence, providing a relatively high-resolution [36] benchmark for supervised R-loop prediction. Following the original DeepER data split, positive sequences were divided into training, validation, and testing sets at a ratio of 7:2:1, with matched negative sequences of equal size in each split. Details of model training, evaluation, and testing are provided in Section 2.2.

### 3.2 The DeepER R-ChIP benchmark contains strong sequence-level signals that are captured by most representations

To better characterize the R-ChIP dataset used in the DeepER study, we first performed motif enrichment analysis on the testing sequences (Fig. S1). The enriched motifs were strongly G-rich and C-rich, consistent with previous empirical observations [17] that canonical R-loop-forming sequences are often associated with GC skew, G-richness, and G-cluster-related sequence patterns. These results suggest that this dataset contains strong sequence-level features that may be detectable by both simple and complex sequence representations.

As an initial proof of concept, we visualized representations from classical encoding methods and pretrained DNA foundation models using UMAP projections of all testing sequences, including 2,889 positive and 2,889 negative sequences (Fig. 2). Across the evaluated representations, positive and negative sequences formed clearly separated groups. This indicates that classical sequence encodings and pretrained DNA foundation model embeddings can capture major differences between R-loop and matched negative sequences in this benchmark, even before applying a downstream classifier.

**Fig. 2:**
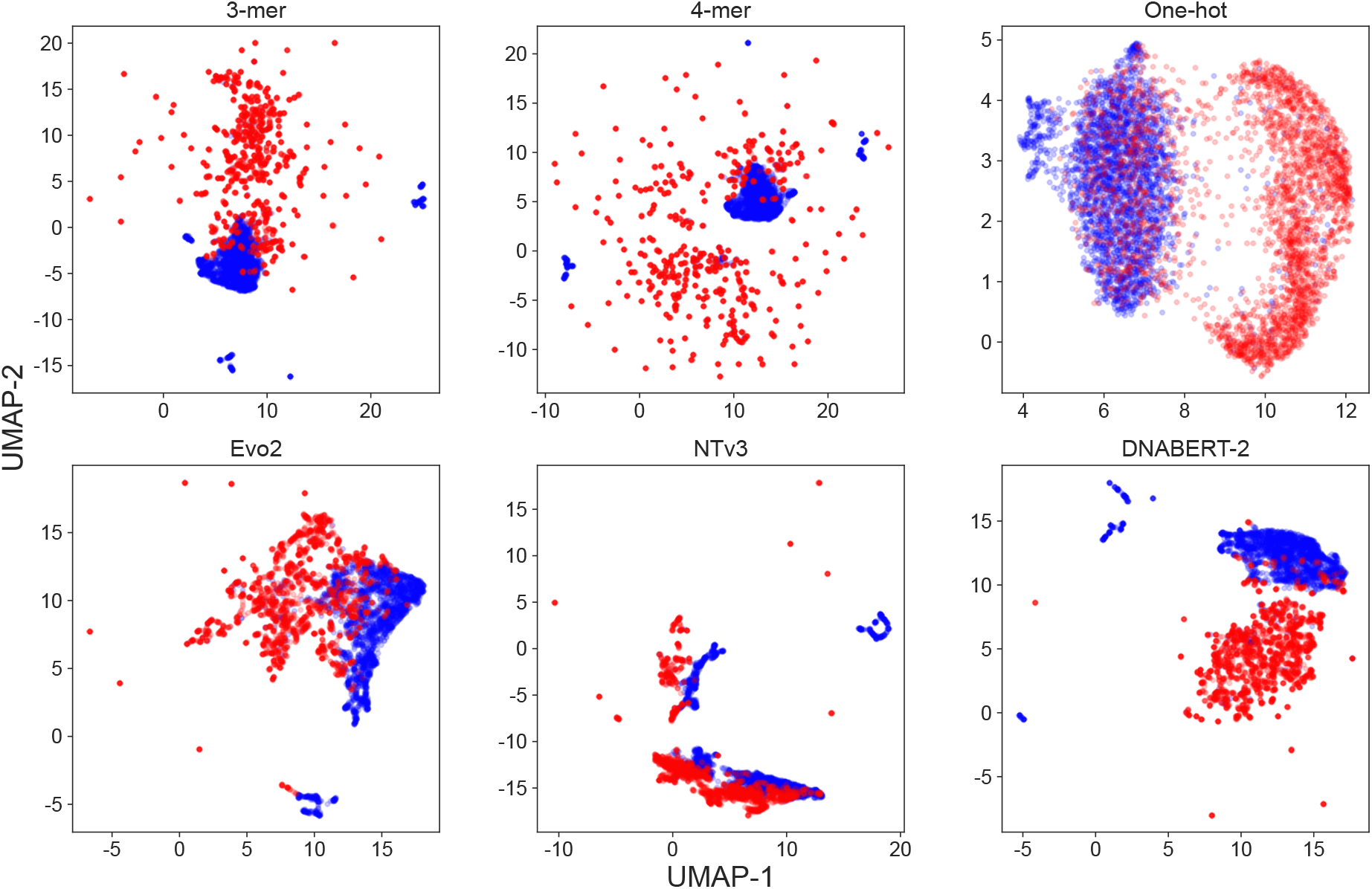
UMAP visualization of testing-set representations from representative DNA sequence representation methods. UMAP projections are shown for 3-mer, 4-mer, one-hot, Evo2, NTv3, and DNABERT-2 representations on the DeepER R-ChIP testing set from HEK293 and K562 cells. For each representation, positive and negative embeddings were combined and projected jointly into two dimensions using a separate UMAP fit with cosine distance. Positive R-loop sequences are shown in red, and matched negative sequences are shown in blue. The testing set contains 2,889 positive and 2,889 negative sequences.

Consistent with this representation-level separation, most machine-learning-based methods achieved strong classification performance on the DeepER R-ChIP testing split (Table 2). The 3-mer and 4-mer baselines performed particularly well, indicating that local sequence composition alone captures a large fraction of the discriminative signal in this dataset. The one-hot baseline also achieved strong performance, suggesting that position-wise sequence information provides additional predictive value for full-length R-loop sequence classification. Pretrained DNA foundation model embeddings, including Evo2, NTv3, and DNABERT-2, also showed high performance, demonstrating that large-scale genomic pretraining can encode R-loop-associated sequence differences in this in-distribution benchmark.

**Table 2:**
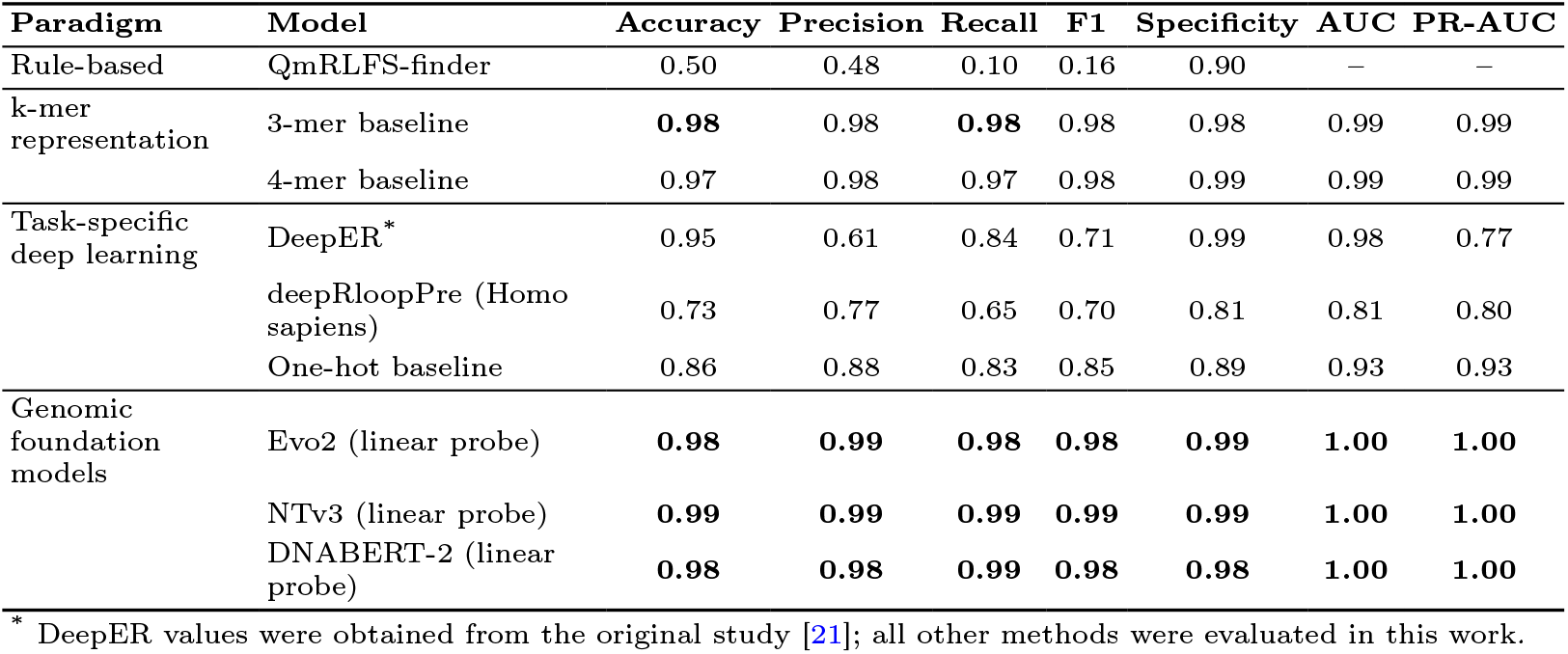
Benchmark comparison of representative R-loop prediction methods on the R-ChIP dataset from HEK293 and K562 cells used in the DeepER study. Following the original DeepER data split, the dataset contains 20,178 positive sequences for training, 5,769 for validation, and 2,889 for testing, with matched negative sequences of equal size in each split. The results reported here are based on the testing set. The top two results for each metric are shown in **bold**.

In contrast, the rule-based QmRLFS-finder showed limited recall despite maintaining high specificity. One possible explanation is that QmRLFS-finder requires sequences to satisfy a complete, empirically defined R-loop-forming pattern, comprising an R-loop initiation zone, a linker region, and an R-loop elongation zone. However, the R-ChIP benchmark dataset was generated by randomly selecting multiple 5-kb intervals within each R-loop region as a data augmentation strategy to improve robustness to R-loop positional variation [21]. As a result, many positive sequences may contain only partial R-loop-associated signals rather than the complete canonical pattern required by QmRLFS-finder. In this setting, QmRLFS-finder would be expected to classify many true R-loop-positive sequences as negative, resulting in high specificity but low recall.

The two task-specific deep learning models also did not show clear improvement over the one-hot baseline. This may partly reflect differences in the effective input sequence length used by each method. In our one-hot baseline, the full 5-kb testing sequence was directly encoded, whereas DeepER and deepRloopPre use shorter sequence windows as model input. Therefore, models that directly encode the full-length sequence may have an advantage in capturing global sequence composition, positional context, and long-range sequence patterns that are relevant for this sequence-level classification task.

In summary, these results show that the DeepER R-ChIP benchmark contains significant, visually separable sequence-level differences between positive and matched negative examples. Both simple sequence encodings and pretrained DNA foundation model embeddings can capture these differences effectively. However, the strong performance observed on this in-distribution benchmark should be interpreted with caution, as it may partly reflect dataset-specific contrasts in sequence composition rather than robust generalization across independent R-loop datasets (see Section *Sequence-level properties help explain dataset-dependent benchmark behavior* for details).

### 3.3 Independent DRIPc-seq validation challenges the generalizability of complex sequence representations

Because most methods performed well on the original DeepER R-ChIP dataset, we next asked whether this performance generalized to an independent DRIPc-seq validation dataset that was previously used to evaluate QmRLFS-finder and R-loop Tracker. This dataset provides a useful cross-platform benchmark because it was generated using a different R-loop mapping strategy. Whereas R-ChIP detects R-loops through catalytically inactive RNase H1 binding [37], DRIPc-seq uses the S9.6 antibody [38] to enrich RNA–DNA hybrids. In addition, none of the evaluated machine-learning-based methods were directly trained on this dataset. Therefore, this dataset serves as an independent blind-test setting for assessing cross-platform generalizability.

The original evaluation of QmRLFS-finder and R-loop Tracker on this dataset was performed at the gene level rather than at the exact signal-segment level. That is, if a gene contained experimentally identified R-loop signals, a prediction was considered successful as long as the algorithm detected at least one R-loop-forming sequence within the same gene, regardless of whether the predicted region matched the exact experimentally identified DRIPc-seq signal. For a more stringent and fair comparison across all methods, we evaluated predictions directly on the exact DRIPc-seq signal regions. We further extracted adjacent upstream or downstream negative control regions for these signals (Section S1.1.2). In total, the validation dataset contained 338 sequences, including 170 positive sequences from validated R-loop regions across 14 genes and 168 adjacent negative control sequences. Compared with the DeepER R-ChIP testing split, this benchmark likely poses a more stringent evaluation, as the negative sequences were sampled from neighboring genomic regions and may therefore be more compositionally similar to positive R-loop sequences. Consistent with this, PCA analysis showed that positive and negative samples were not clearly separated (Fig. S2). Motif enrichment analysis of this dataset revealed that the top enriched motifs (Fig. S3) differed substantially from those identified in the DeepER R-ChIP testing set (Fig. S1). Although G-cluster-like patterns were still observed, the enriched motifs also included A- and T-rich components, potentially reflecting linker-like sequence contexts, gene-specific composition, or other noncanonical features of experimentally detected R-loops. Together, these results suggest that the two benchmarks differ in sequence composition and that R-loop-associated signals cannot be reduced simply to G- or C-richness. This finding suggests that the sequence features captured by QmRLFS-style rules, while biologically informative, may not fully explain R-loop-associated sequence signatures across different benchmark designs.

Overall, performance on this independent validation set was modest across all methods (Table 3). Unexpectedly, the one-hot baseline achieved the best overall balance, with the highest accuracy, recall, and F1 score among the tested methods. Evo2 also showed competitive performance. This pattern may reflect the importance of capturing sequence features across longer genomic intervals, rather than relying only on short local motifs. The 3-mer and 4-mer baselines, together with NTv3 and DNABERT-2, showed slightly lower balanced classification performance, but they produced the highest or near-highest AUC and PR-AUC values among the tested methods. This suggests that simple sequence-composition features retain sufficient information to distinguish R-loop signals from adjacent negative regions.

**Table 3:**
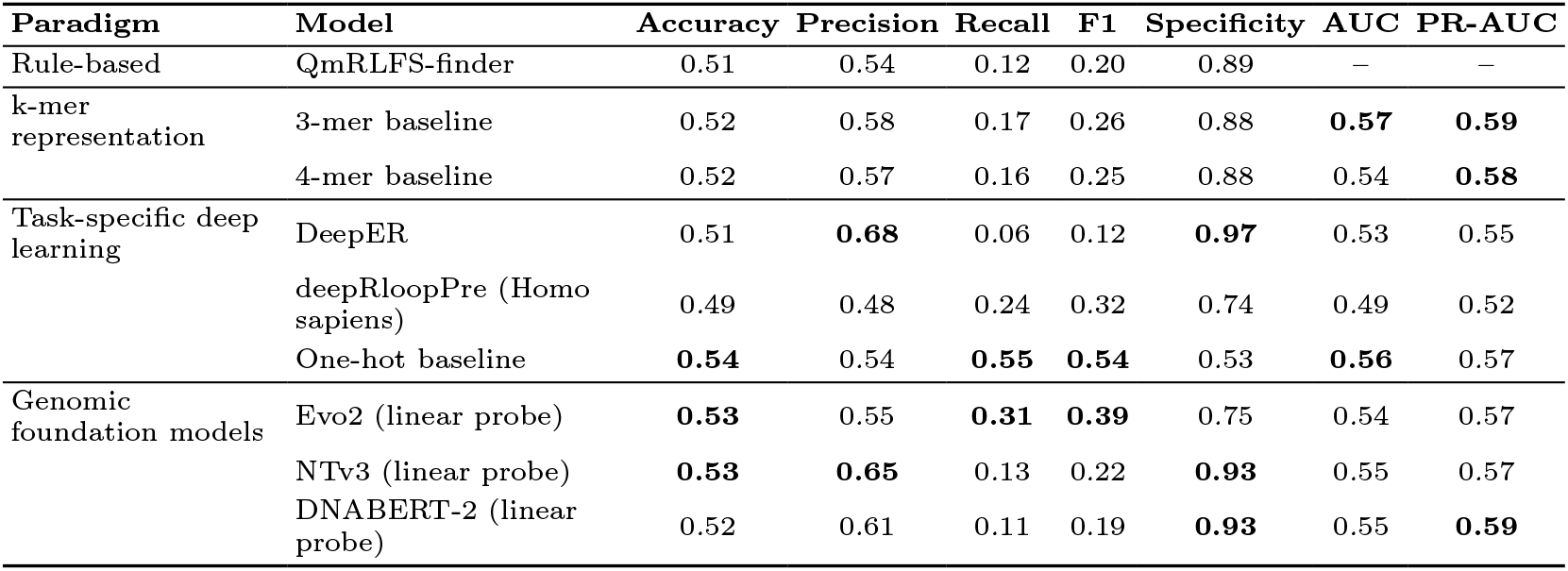
Benchmark comparison of representative R-loop prediction methods on a DRIPc-seq validation dataset of 338 sequences, including 170 positive sequences from validated R-loop regions in 14 genes and 168 negative control sequences sampled from the corresponding upstream or downstream neighboring regions. This testing dataset was originally used in the QmRLFS-finder and the R-loop Tracker studies. The top two results for each metric are shown in **bold**.

In contrast, the task-specific deep learning models showed uneven behavior. DeepER achieved the highest precision and specificity, with values of 0.68 and 0.97, respectively, but its recall was only 0.06. This means that DeepER made very few false-positive predictions but missed most true R-loop regions, indicating highly conservative predictions and limited sensitivity to previously unseen R-loop signals in this cross-platform benchmark. deepRloopPre showed higher recall than DeepER on the DRIPc-seq validation set, despite its weaker performance on the DeepER R-ChIP benchmark. This difference may reflect platform similarity, as deepRloopPre was trained on ssDRIP-seq [39] datasets and both ssDRIP-seq and DRIPc-seq rely on S9.6 antibody-based enrichment of RNA–DNA hybrids. Thus, deepRloopPre may better capture sequence features associated with antibody-based DRIP-family signals than those represented in the R-ChIP benchmark. However, its overall accuracy, F1 score, AUC, and PR-AUC remained limited, indicating that improved recall alone did not translate into strong balanced prediction performance.

Taken together, the divergence between Table 2 and Table 3 supports the central observation of this study: DNA foundation model embeddings can perform extremely well in an in-distribution R-ChIP benchmark, but this advantage does not consistently transfer to an independent, unseen, cross-platform validation setting. These results suggest that R-loop prediction performance is strongly influenced by dataset construction, negative-control selection, experimental platform, and sequence-composition differences between positive and negative regions.

### 3.4 Higher-confidence R-loopBase sequences are easier to detect

The results above highlight the inherent difficulty of establishing robust R-loop classification benchmarks, due to cross-platform heterogeneity [40] and uncertainty in defining reliable negative examples. Although positive R-loop regions can be supported by experimental enrichment, the absence of a detected signal does not necessarily indicate that a sequence lacks R-loop-forming potential. Unannotated or signal-depleted regions may instead reflect limited assay sensitivity, cell-type- or condition-specific R-loop formation, or platform-dependent detection biases, introducing label uncertainty into negative sets. We therefore shifted our subsequent analysis from binary classification to recall-based evaluation, which measures how well each method recovers experimentally identified R-loop regions without requiring a predefined negative set.

For this analysis, we selected R-loopBase [31] as a multi-platform reference of human R-loop zones (Section S1.1.3). R-loopBase provides high-confidence R-loop annotations by integrating 107 high-quality genome-wide R-loop mapping datasets generated using 11 different technologies. R-loop zones are assigned consensus levels according to the number of technologies by which they are detected. For example, Level 4 zones are detected by at least four technologies. Thus, higher consensus levels represent increasingly well-supported R-loop annotations, with Level 9 containing the smallest but highest-confidence set of R-loop zones. We evaluated model recall across R-loopBase consensus levels from Level 4 to Level 9 (Fig. 3). This range was chosen because Levels 1–3 contain many fragmented and low-confidence regions, whereas Levels 4–9 retain sufficient testing sequences while representing increasingly confident R-loop annotations.

**Fig. 3:**
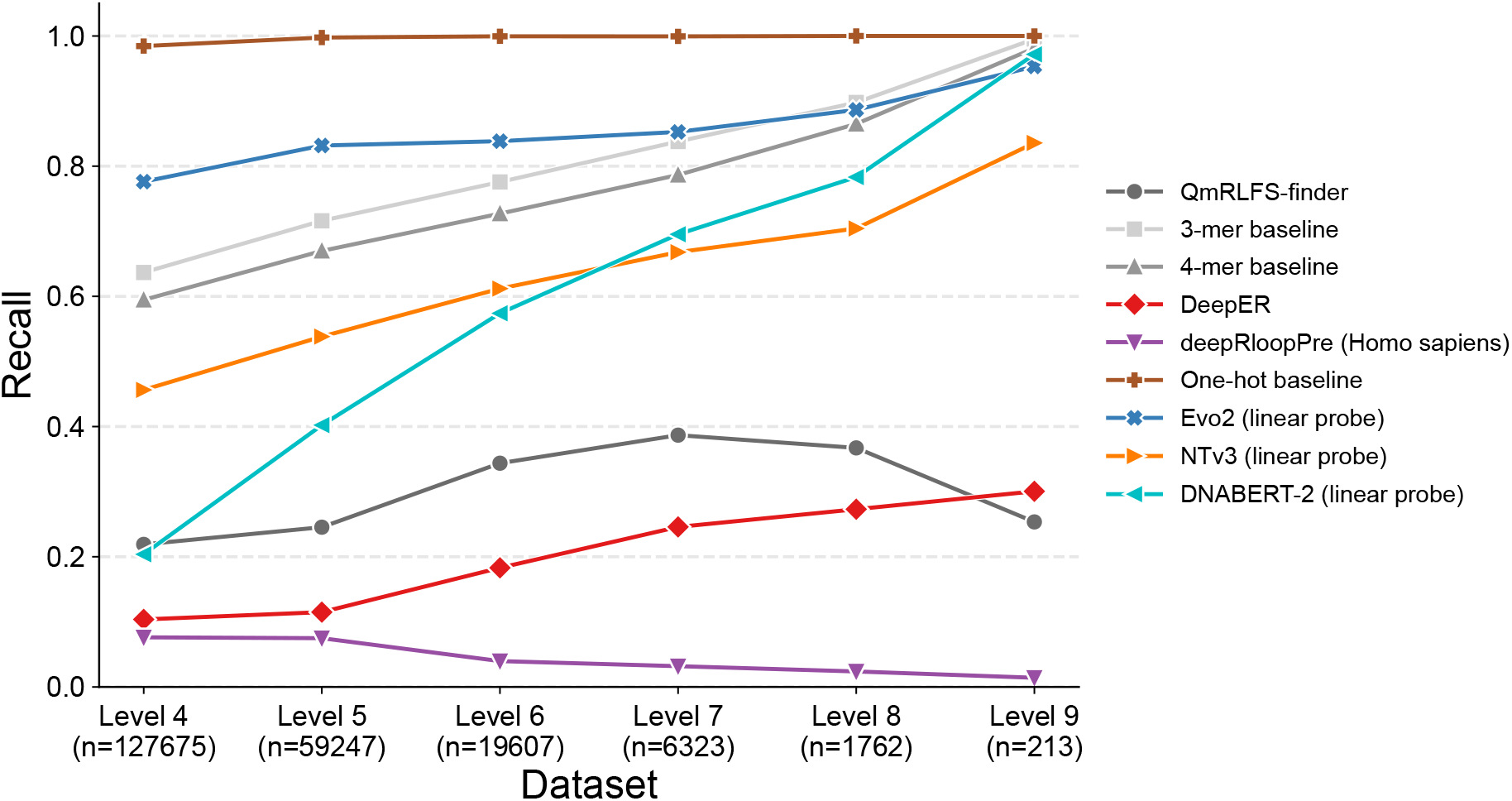
Comparison of model recall across R-loopBase consensus levels. Recall was evaluated for QmRLFS-finder, classical sequence encodings, task-specific deep learning models, and DNA foundation model embeddings using linear probes on R-loopBase regions spanning consensus levels 4 to 9. Each point represents the fraction of sequences at a given consensus level that were correctly identified as R-loop-forming by the corresponding method. The number of testing sequences (*n*) at each level is shown below the corresponding x-axis label.

For most methods, recall increased with increasing R-loopBase consensus level. This trend suggests that higher-confidence R-loop regions contain stronger or more canonical sequence signals that are easier for sequence-based models to detect. The one-hot baseline maintained high recall across all R-loopBase levels, reaching near-complete recall at the highest consensus levels. Evo2 ranked second overall, suggesting that its long-context genomic representation captures substantial R-loop-associated sequence information. The k-mer baselines also showed steadily increasing recall as consensus levels rose, indicating that compositional signals become more pronounced in high-confidence R-loop regions. NTv3 and DNABERT-2 improved substantially from Level 4 to Level 9, suggesting that foundation models can recover canonical R-loop-associated sequence signals more effectively. By comparison, QmRLFS-finder showed only modest recall across all consensus levels, recovering only approximately one-third of Level 9 regions. This pattern indicates that its predefined RIZ–linker–REZ rule captures only a subset of high-fidelity consensus R-loop regions.

Among task-specific deep learning models, recall patterns are more variable. DeepER shows only a modest increase in recall across R-loopBase consensus levels and consistently underperforms compared with most classical and foundation-model-based approaches. deepRloopPre shows comparatively lower recall across all tested consensus levels. Because DeepER and deepRloopPre use broadly similar one-hot-based neural network architectures (Table 1), the observed difference in recall is likely driven by differences in training data rather than architecture alone. In particular, deepRloopPre was trained on ssDRIP-seq-derived datasets, which generally provide lower-resolution R-loop mapping than R-ChIP [36]. This may reduce its ability to identify the more precisely defined and multi-platform-supported R-loop zones represented in R-loopBase.

Overall, the R-loopBase benchmark supports two main conclusions. First, most machine-learning-based methods are more successful at recovering higher-confidence R-loop zones, implying that these regions are enriched for stronger sequence-level features associated with R-loop formation. Second, simple representation methods and advanced foundation model embeddings can both capture these signals, and increased model complexity does not guarantee better recall.

### 3.5 Cross-species evaluation reveals different generalization patterns across representation paradigms

Previous R-loop prediction tools have largely been developed and evaluated in target-specific settings, with some methods focused primarily on the human genome, such as QmRLFS-finder and DeepER, and others designed for plant species, such as deepRloopPre. In contrast, DNA foundation models are pretrained on large-scale multi-species genomic corpora and have shown promise in cross-species genome annotation tasks [13,5], even without extensive task-specific post-training. We therefore evaluated cross-species generalization of these models on genome stability-related tasks such as R-loop prediction.

To evaluate cross-species generalization, we curated R-loop datasets from multiple organisms (Section S1.1.4). The number of R-loop regions varied substantially across species, likely reflecting differences in experimental resolution, detection sensitivity, genome size, and biological context. Notably, *Z. mays* contained many more R-loop regions than the other species, possibly due in part to its large genome and abundant pericentromeric repeats [41]. This distinct genomic context suggests that *Z. mays* may pose a more challenging setting for evaluating cross-species R-loop prediction.

The cross-species benchmark revealed substantial variability across both datasets and model classes (Fig. 4). Consistent with the human benchmark results (Tables 2 and 3), the one-hot baseline shows the strongest overall recall across species, suggesting that position-specific sequence representations remain highly effective for R-loop detection. Classical k-mer baselines also perform well in several plant species and *D. melanogaster* ; however, their recall declined in species such as *D. rerio* and *M. musculus*, which may reflect more divergent and complex genomic backgrounds. These results suggest that sequence-composition features alone, without preserving positional or contextual organization, may limit the cross-organism transferability of predictive models.

**Fig. 4:**
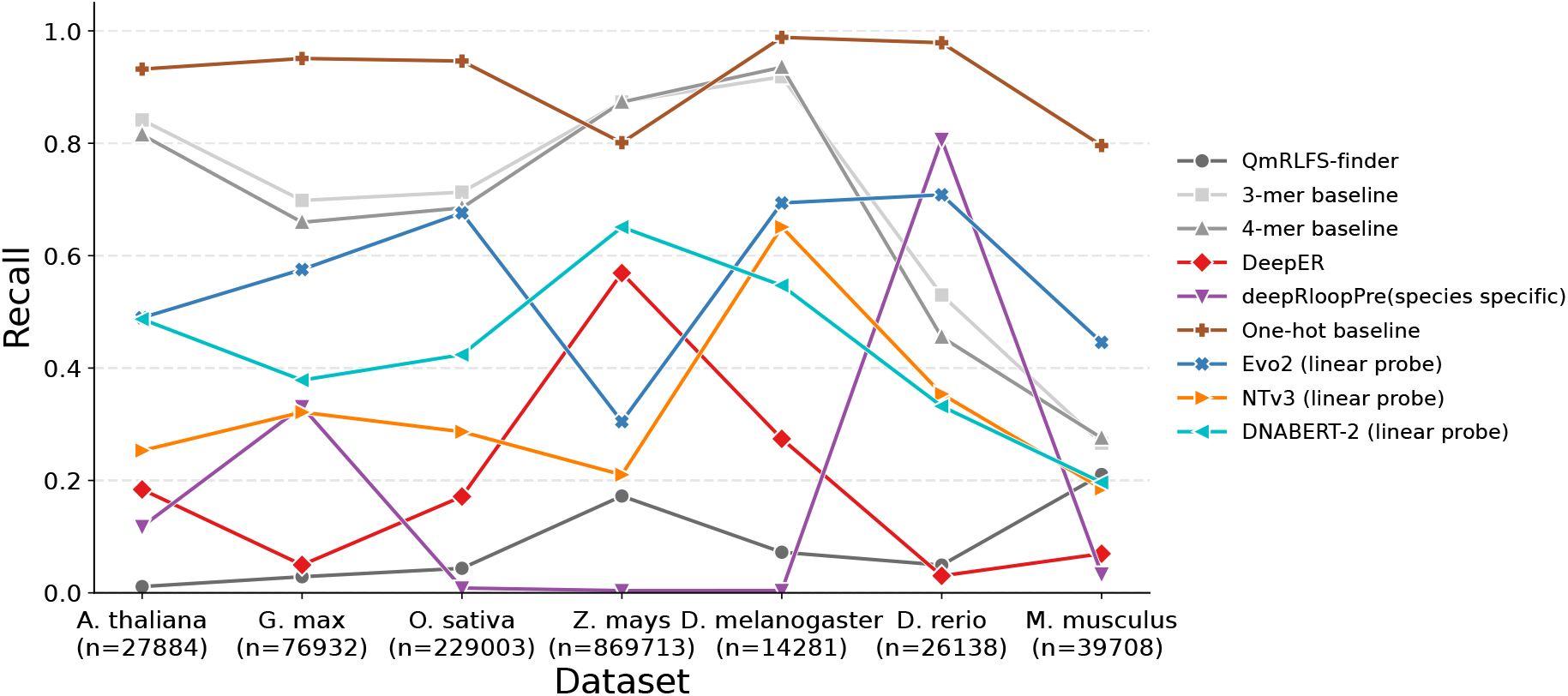
Comparison of recall across different methods on the cross-species R-loop datasets curated in this study. For deepRloopPre, the corresponding species-specific model was used when available, including models for *Arabidopsis thaliana, Danio rerio*, and *Oryza sativa*. Otherwise, the human model was applied. as *D. rerio*, when a matching model was available, but these improvements did not translate into strong performance across the full benchmark.

Foundation models showed mixed performance. Evo2 achieved a relatively stable recall across several species, except in *Z. mays*, and was often comparable to k-mer baselines. By contrast, NTv3 and DNABERT-2 generally showed more modest recall. These results suggest that pretrained DNA language models can capture some transferable sequence features relevant to R-loop prediction, but sequence-only pretraining may not be sufficient to fully model species-specific R-loop signals. The stronger performance of Evo2 may also reflect scaling effects [42], as Evo2 has substantially more parameters than NTv3 and DNABERT-2.

Task-specific deep learning models still showed limited transferability. DeepER recovered only a subset of R-loop regions outside its original human-centered setting, with detectable recall mainly in *Z. mays*. For deepRloopPre, species-specific pretrained models provided recall gains in selected organisms, such Together, these benchmarks show that cross-species R-loop prediction remains challenging. In this setting, simple supervised representations that preserve sequence and positional information generalized surprisingly well, whereas both general-purpose foundation models and task-specific deep learning models showed species-dependent limitations.

### 3.6 Sequence-level properties help explain dataset-dependent benchmark behavior

Previous studies have shown that R-loop-forming sequences are associated with distinct G-rich and compositionally biased sequence features [17,19,43,35,18,44], including G-clusters, G-quadruplex-forming motifs, GC skew, and CpG islands. However, R-loop sequence composition is complex, heterogeneous, and incompletely characterized, especially across different species and experimental platforms. To provide a more interpretable explanation for the benchmark results, we summarized multiple sequence-level properties across the evaluated R-loop datasets, including GC content, G content, GC skew, AT skew, CpG density, G-run density, sequence entropy, and G4 motif fraction (Section 2.3; Table 4). These properties differed substantially across datasets, indicating that the R-loop benchmarks evaluated in this study represent diverse sequence contexts.

**Table 4:** Sequence-level properties summarized across R-loop datasets.

| Dataset | Subset | *N* | GC (%) | G (%) | GC Skew | AT Skew | CpG (%) | G-run<sup>a</sup> | Entropy | G4 (%) |
| --- | --- | --- | --- | --- | --- | --- | --- | --- | --- | --- |
| **DeepER R-ChIP** |  |  |  |  |  |  |  |  |  |  |
|  | Positive Training | 20178 | 49.9 | 25.3 | 0.016 | −0.007 | 3.4 | 16.9 | 0.994 | 31.4 |
|  | Evaluation | 5769 | 49.6 | 25.2 | 0.016 | −0.007 | 3.4 | 16.8 | 0.994 | 29.8 |
|  | Testing | 2889 | 49.8 | 25.2 | 0.014 | −0.005 | 3.4 | 16.7 | 0.994 | 29.8 |
|  | Negative Training | 20178 | 39.0 | 19.3 | −0.011 | −0.029 | 0.8 | 8.2 | 0.976 | 4.5 |
|  | Evaluation | 5769 | 39.0 | 19.3 | −0.011 | −0.029 | 0.8 | 8.2 | 0.976 | 4.5 |
|  | Testing | 2889 | 38.8 | 19.2 | −0.012 | −0.029 | 0.8 | 8.1 | 0.976 | 5.1 |
| **R-loop Tracker (DRIPc-seq)** |  |  |  |  |  |  |  |  |  |  |
|  | Positive R-loop | 170 | 41.9 | 22.4 | 0.067 | 0.014 | 1.1 | 12.0 | 0.978 | 4.7 |
|  | Adjacent Negative | 168 | 40.4 | 20.6 | 0.020 | −0.006 | 0.9 | 10.1 | 0.979 | 5.4 |
| **R-loopBase (Consensus Levels)** |  |  |  |  |  |  |  |  |  |  |
|  | Level 4 | 127675 | 53.4 | 28.5 | 0.072 | 0.040 | 3.0 | 20.4 | 0.974 | 5.2 |
|  | Level 5 | 59247 | 56.6 | 30.4 | 0.080 | 0.049 | 3.6 | 23.3 | 0.970 | 5.4 |
|  | Level 6 | 19607 | 61.3 | 33.2 | 0.085 | 0.037 | 4.9 | 28.7 | 0.959 | 6.5 |
|  | Level 7 | 6323 | 64.7 | 35.4 | 0.094 | 0.038 | 6.2 | 32.3 | 0.947 | 6.2 |
|  | Level 8 | 1762 | 66.2 | 37.1 | 0.121 | 0.054 | 7.3 | 34.4 | 0.939 | 5.5 |
|  | Level 9 | 213 | 68.1 | 40.5 | 0.188 | 0.155 | 9.5 | 35.5 | 0.921 | 3.3 |
| **Cross-species Curated Set** |  |  |  |  |  |  |  |  |  |  |
|  | *A. thaliana* | 27884 | 39.2 | 23.1 | 0.175 | 0.126 | 2.9 | 7.3 | 0.962 | 0.2 |
|  | *G. max* | 76932 | 36.0 | 22.9 | 0.263 | 0.200 | 1.9 | 9.2 | 0.919 | 0.6 |
|  | *O. sativa* | 229003 | 43.6 | 21.8 | 0.000<sup>b</sup> | −0.001 | 4.2 | 8.5 | 0.958 | 0.4 |
|  | *Z. mays* | 869713 | 46.9 | 23.4 | 0.000<sup>c</sup> | 0.000<sup>d</sup> | 4.4 | 10.7 | 0.983 | 1.6 |
|  | *D. melanogaster* | 14281 | 50.2 | 21.3 | −0.154 | 0.006 | 5.3 | 6.9 | 0.984 | 0.4 |
|  | *D. rerio* | 26138 | 41.5 | 29.1 | 0.404 | 0.355 | 1.9 | 12.6 | 0.889 | 0.8 |
|  | *M. musculus* | 39708 | 47.7 | 23.9 | 0.001 | 0.000<sup>e</sup> | 1.6 | 13.4 | 0.974 | 6.8 |

<sup>a</sup> G-run density is defined as the number of $G_{\geq 3}$ runs per kb.
<sup>b-e</sup> Precise values for entries rounded to 0.000: *O. sativa* GC skew = $4.04 \times 10^{-4}$ ; *Z. mays* GC skew = $7.54 \times 10^{-5}$ , AT skew = $-3.19 \times 10^{-4}$ ; *M. musculus* AT skew = $-2.25 \times 10^{-4}$ .
*Note:* *N* denotes the total number of sequences in each subset. GC content, G content, CpG density, and G4 motif fraction are reported as percentages (%). GC skew, AT skew, and normalized Shannon entropy are reported as dataset-level mean values. Values shown as 0.000 reflect rounding to three decimal places and should not be interpreted as exact zeros.

In the DeepER R-ChIP dataset, positive sequences showed substantially higher GC content, G content, G-run density, and G4 motif fraction than matched negative sequences. This strong compositional contrast likely explains why most DNA representations performed well on the DeepER testing set (Table 2) and why positive and negative samples were clearly separated in the UMAP projection (Fig. 2). By comparison, the DRIPc-seq validation set showed much smaller compositional differences between positive R-loop sequences and adjacent negative sequences, particularly for C/G-related features. This is consistent with the PCA analysis, where positive and negative samples were not clearly separated (Fig. S2), and likely contributes to the reduced performance observed across methods (Table 3). Notably, positive sequences in this dataset showed elevated AT skew, suggesting that these validated R-loop regions may not be driven solely by canonical G-rich or GC-skewed sequence features. This observation is also compatible with previous reports linking R-loop formation to transcription termination and poly(A)-associated regions [15], although additional context-dependent factors are presumably involved.

R-loopBase consensus levels showed a clear increase in GC content, G content, GC skew, CpG density, and G-run density from Level 4 to Level 9. This compositional strengthening provides a plausible explanation for the increased recall (Fig. 3) observed at higher consensus levels for many methods. In other words, higher-confidence R-loop regions are enriched for more canonical G-rich and GC-skewed sequence features, making them easier to recover using both classical and learning-based representations.

The cross-species datasets also displayed broad compositional variation, reflecting the diverse sequence contexts in which R-loops form across organisms. For example, GC content ranged from relatively low values in *G. max* and *A. thaliana* to higher values in *D. melanogaster* and *M. musculus*. GC skew and AT skew also varied substantially across species. *O. sativa, Z. mays*, and *M. musculus* showed weaker G-C or A-T strand asymmetry, whereas *G. max* and *D. rerio* showed stronger skew patterns. In addition, *D. rerio* showed markedly low sequence entropy, suggesting lower local sequence complexity in the evaluated R-loop regions, consistent with the repetitive features of the zebrafish genome [45]. These differences provide a plausible explanation for the inconsistent cross-species recall patterns observed in Fig. 4.

Collectively, these analyses indicate that R-loop-associated sequence features vary substantially across datasets, platforms, and species. Sequence-based classifiers trained under one compositional regime may therefore show reduced transferability to another, especially when the target species differs in GC content, strand asymmetry, repetitive sequence structure, or G4-associated features. More broadly, these results suggest that R-loop prediction cannot be fully explained by a single universal sequence grammar [20]. Current sequence-only foundation models capture some transferable sequence patterns, but may still miss determinants of R-loop formation shaped by assay conditions, species-specific sequence composition, chromatin state, and broader epigenomic context [16].

## 4 Discussion

In this study, we systematically benchmarked classical and foundation model-based DNA sequence representations for R-loop-forming sequence prediction, a genome stability-associated task that remains underexplored in current DNA foundation model evaluations [13,14,23]. To focus the comparison on representation quality rather than downstream classifier complexity, we evaluated sequence representations using a unified linear-probe framework. A central observation is that strong performance on a dataset with clear positive–negative compositional contrast did not consistently transfer to independent benchmarks generated from different experimental platforms or biological contexts. Simple sequence representations, including *k*-mer and one-hot encodings, remained highly competitive across evaluations. In particular, the one-hot baseline, which directly preserves sequence and positional information across 5-kb windows, showed strong performance in recovering both high-confidence human R-loop regions and cross-species R-loop signals. These results suggest that much of the recoverable signal in current R-loop benchmarks is still driven by sequence-level features, and that simple baselines are essential controls when evaluating more complex models.

Accurate and generalizable prediction of R-loop-forming sequences remains challenging for two main reasons. First, R-loop biology is intrinsically context-dependent [20,22,16]. R-loop distribution and sequence composition vary across species, genomic regions, and regulatory environments. In mammals and several plant species, such as *A. thaliana* and *O. sativa*, R-loops are often enriched in transcriptionally active regions, including promoters and termination regions [15,26,39,46]. In contrast, R-loops in *Z. mays* have been reported near centromeric regions [41], whereas R-loops in *D. melanogaster* can be associated with Polycomb response elements [47]. These examples indicate that R-loop formation cannot be fully explained by a single universal sequence grammar [20]. Future models may therefore need to incorporate broader genomic and epigenomic context, including transcriptional activity [31,48,49,50], chromatin accessibility [51,52,53], histone modifications [54], DNA methylation [55,56,57], and other regulatory profiles.

Second, defining true negative R-loop-forming sequences remains difficult. The absence of an observed R-loop signal in a given assay, cell type, or condition does not necessarily indicate that a genomic region lacks R-loop-forming potential, because the same region may exhibit detectable signals in a different biological context [36]. This uncertainty makes negative-control construction a major source of variability in R-loop classification benchmarks. In the DeepER R-ChIP dataset, positive and negative sequences showed clear compositional separation, suggesting that models could partly distinguish the two classes using broad sequence-composition features. By contrast, the DRIPc-seq validation set used adjacent genomic regions as negative controls, making the negative sequences more compositionally similar to positive R-loop regions and therefore harder to separate. This adjacent-control design poses a more stringent benchmark, as it requires finer discrimination of R-loop-associated signals within similar local genomic contexts rather than reliance on broad compositional differences. Together, these observations suggest that benchmarks based on randomly sampled negative regions may overestimate model generalizability when positive and negative sets differ significantly in general sequence composition. Although the strong recall of the one-hot baseline indicates high sensitivity to R-loop-forming sequences, its apparent advantage may be reduced in more stringent positive-versus-native-background tasks that require discrimination from compositionally similar genomic regions. Future R-loop benchmarks should therefore incorporate more carefully matched negative sets, cross-platform validation, and broader celltype coverage to better assess whether models capture generalizable R-loop-forming sequence features rather than dataset-specific biases.

Our results also highlight both the utility and limitations of pretrained DNA foundation models [6]. These models captured useful R-loop-associated sequence signals, especially in the in-distribution R-ChIP benchmark and high-confidence R-loopBase regions. However, their advantages were less consistent in independent DRIPc-seq and cross-species evaluations, where classical encodings often remained competitive. This suggests that sequence-only DNA foundation models may not fully capture genome stability-associated tasks whose determinants depend on assay conditions, species-specific sequence composition, chromatin state, and broader epigenomic context. For R-loop prediction and related genomic tasks, future foundation models may benefit from integrating DNA sequence with multi-omic regulatory information [58,59] rather than relying on sequence pretraining alone. Overall, our benchmark supports R-loop prediction as a complementary genome stability-associated test case for evaluating whether DNA foundation models generalize beyond conventional gene regulatory prediction tasks.

## Acknowledgments

This work was supported by startup funding from North Carolina State University and National Science Foundation (NSF) Grant MCB-2540505, with additional support from the NC State Genetics and Genomics Academy and the Comparative Medicine Institute. We are grateful to Dr. Fangpu Han (corresponding author of PMID: 34244230) and his lab members for sharing the processed maize R-loop data with us. We also appreciate the fruitful discussions with Dr. Jianghao Shen and Dr. Chun-Ying Lee. All trained R-loop classifiers developed in this work are available at https://github.com/LinResearchGroup-NCSU/RLoopBench.

