## Supplementary Materials for "R-loop Prediction Reveals Generalization Limits of DNA Foundation Models Beyond Regulatory Genomics"

### S1 Supplementary Texts

#### S1.1 Data Curation

This study used four R-loop datasets: 1) the R-ChIP dataset originally generated in the DeepER study [21]; 2) the DRIPc-seq dataset used by QmRLFS-finder [25] and R-loop Tracker [30]; 3) Level 4 to Level 9 R-loopBase datasets [31]; and 4) cross-species R-loop datasets curated in this study. Below, we briefly describe each dataset and the necessary preprocessing steps.

**S1.1.1 R-ChIP Dataset from the DeepER Study** We used the processed R-ChIP dataset released by the DeepER study, directly downloaded from its GitHub repository (`https://github.com/NjuChenlab/DeepER/tree/main/data`). The dataset was curated from R-ChIP peaks conserved between K562 and HEK293 cells [60] and further supported by independent R-loop mapping datasets from R-loopBase. Positive samples were 5-kb augmented intervals containing curated R-loop regions, while negative samples were sampled 5-kb genomic intervals without overlap with known R-loop regions. The training, validation, and testing sets contained 20,178, 5,769, and 2,889 positive sequences, respectively, with an equal number of negative sequences in each split. We treated R-loop-forming sequences as positive samples and R-loop-negative sequences as negative samples for binary classification in Section 2.2.2.

**S1.1.2 DRIPc-seq Dataset from R-loop Tracker** We used the DRIPc-seq [38] validation dataset originally used for QmRLFS-finder and its web-based implementation, R-loop Tracker [30]. The dataset was obtained from the public repository (`https://github.com/jan-havlik/validation_dataset`). It contains strand-specific R-loop regions across 14 genes, with genomic coordinates based on GRCh37/hg19. In total, 170 positive R-loop regions were included.

To construct a binary classification dataset, we generated matched negative sequences for these positive R-loop regions. For each positive region, one negative region with the same chromosome, length, and strand was selected from a nearby genomic interval. The upstream adjacent region was prioritized, followed by the downstream adjacent region when the upstream candidate was not valid. A 500-bp buffer was imposed between positive and negative regions, and candidate negative regions overlapping any known positive R-loop region were excluded. If no valid adjacent region was available, a random region from the same chromosome was selected under the same constraints. Sequences containing ambiguous nucleotides were ignored. Using this strategy, we generated matched negative sequences for 168 of the 170 positive R-loop regions. The remaining two regions were excluded because no valid negative region of the same length could be identified on the same chromosome without overlapping positive R-loop regions or exceeding chromosome boundaries.

**S1.1.3 R-loopBase Level 4–Level 9 Datasets** R-loop mapping datasets were obtained from R-loopBase [31] and downloaded from the R-loopBase data portal (https://rloopbase.nju.edu.cn/download.jsp). R-loopBase curates R-loop zones detected by multiple experimental technologies across diverse cell types and genomic contexts. Each R-loop zone is assigned to one of nine confidence levels, with higher levels indicating stronger multi-technology support and therefore higher expected reliability. We used Level 4 to Level 9 R-loop regions as benchmark datasets to evaluate model recall across confidence tiers. The number of regions in each level is summarized in Figure 3.

**S1.1.4 Cross-Species R-loop Datasets Curated in This Study** We curated R-loop datasets from four plant species (*A. thaliana*, *G. max*, *O. sativa*, and *Z. mays*) and three animal species (*D. melanogaster*, *D. rerio*, and *M. musculus*). Across all species, only R-loop regions on nuclear chromosomes were retained, whereas mitochondrial, chloroplast, unplaced, unlocalized, and other non-standard contigs were excluded. We briefly describe the corresponding data sources and preprocessing steps.

The *Arabidopsis thaliana* R-loop data were obtained from GEO accession GSM3214328, which provides wild-type R-loop signals for both forward and reverse strands. The forward-strand signal corresponds to wR-loops, representing R-loop formation with single-stranded DNA on the Watson strand and a DNA:RNA hybrid on the Crick strand, whereas the reverse-strand signal corresponds to cR-loops. The original files were provided as normalized coverage tracks in BigWig format at 1-bp resolution, where each genomic position has a normalized R-loop coverage score. Therefore, we converted these base-level signal tracks into continuous R-loop regions.

Specifically, the forward- and reverse-strand signal tracks were processed separately using a strict region-calling procedure. The original 1-bp-resolution BigWig files were first converted to bedGraph format. Positions with normalized coverage scores $\geq 3.5$ were selected as seed intervals, and directly adjacent or overlapping seed intervals were merged to form candidate R-loop regions. Candidate regions were retained if they showed a mean coverage $> 5.0$ across the region and had a length between 150 bp and 5,000 bp. The retained forward- and reverse-strand regions were assigned to the positive and negative strands, respectively, and merged into the final *A. thaliana* R-loop BED file.

Sequences for the final R-loop regions were extracted from the TAIR10 reference genome downloaded from Ensembl Plants release 75 (ftp://ftp.ensembl.org/pub/release-75/fasta/arabidopsis_thaliana/dna/Arabidopsis_thaliana.TAIR10.dna.toplevel.fa.gz).

R-loop datasets for *Glycine max*, *Oryza sativa*, and *Danio rerio* were obtained from the GEO database. For *Glycine max*, GSM6671425 and GSM6671426 were used; for *Oryza sativa*, GSM4972197 and GSM4972198 were used; and for *Danio rerio*, GSM5557861 was used. The corresponding reference genomes were obtained from GenBank assemblies GCA_003349995.2, GCA_001623365.2, and GCA_000002035.2, respectively. R-loop sequences were extracted following the same procedure used for *Arabidopsis thaliana*, yielding 76,932 R-loop regions for *Glycine max*, 229,003 for *Oryza sativa*, and 26,138 for *Danio rerio*.

R-loop data for *Zea mays* in BED format were directly provided by the authors of [41]. The corresponding reference genome was downloaded from Ensembl Plants release 62: https://ftp.ensemblgenomes.ebi.ac.uk/pub/plants/release-62/fasta/zea_mays/dna/Zea_mays.Zm-B73-REFERENCE-NAM-5.0.dna.toplevel.fa.gz. Sequences shorter than 150 bp were filtered out, resulting in 759,673 R-loop sequences for testing.

R-loop datasets for *Drosophila melanogaster* and *Mus musculus* were obtained from the GEO database. For *Drosophila melanogaster*, R-loop peaks identified in S2 cells from GSE127329 were used, including GSE127329_S2_F.bed.gz and GSE127329_S2_R.bed.gz. The corresponding dm3 reference genome was downloaded from UCSC: https://hgdownload.soe.ucsc.edu/goldenPath/dm3/bigZips/dm3.fa.gz. For *Mus musculus*, GSM2104456 was used with the GenBank genome assembly GCA_000001635.1. This yielded 14,281 and 39,708 R-loop sequences for *Drosophila melanogaster* and *Mus musculus*, respectively.

#### S1.2 Motif discovery using STREME

We used STREME [61] from the MEME Suite to identify candidate sequence motifs enriched in R-loop-forming regions. Positive R-loop sequences were compared against matched negative sequences using DNA mode. Motif widths were restricted to 6–20 bp, and the top three motifs were reported for each analysis. The top three motifs were visualized as sequence logos based on the position probability matrices reported by STREME. For each motif, the motif width, number of contributing sites, and E-value are shown.

### S2 Supplementary Figures

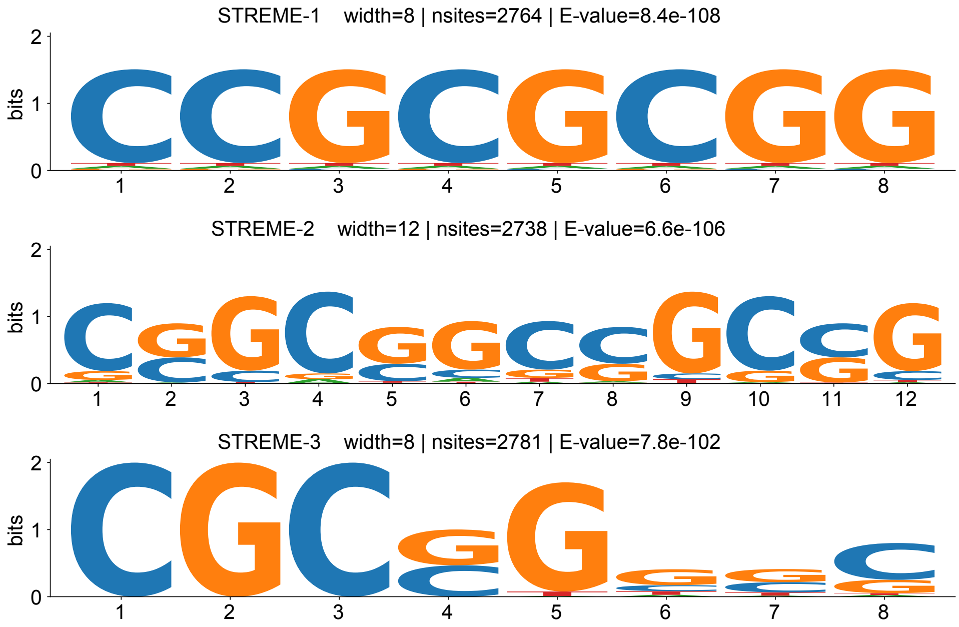


Fig. S1: **Candidate enriched sequence motifs identified from the DeepER R-ChIP testing dataset.** Sequence logos are shown for the top three motifs identified by STREME [61] using the testing portion of the DeepER R-ChIP dataset. Positive R-loop sequences were compared against negative sequences, with 2,889 positive and 2,889 negative sequences included in the analysis. For each motif, STREME reports the motif width, the number of contributing sites used to construct the motif, and the E-value, which represents the expected number of motifs with equal or greater enrichment that would be found by chance after accounting for the multiple candidate motifs and motif widths evaluated during the search. These values are shown above each sequence logo.

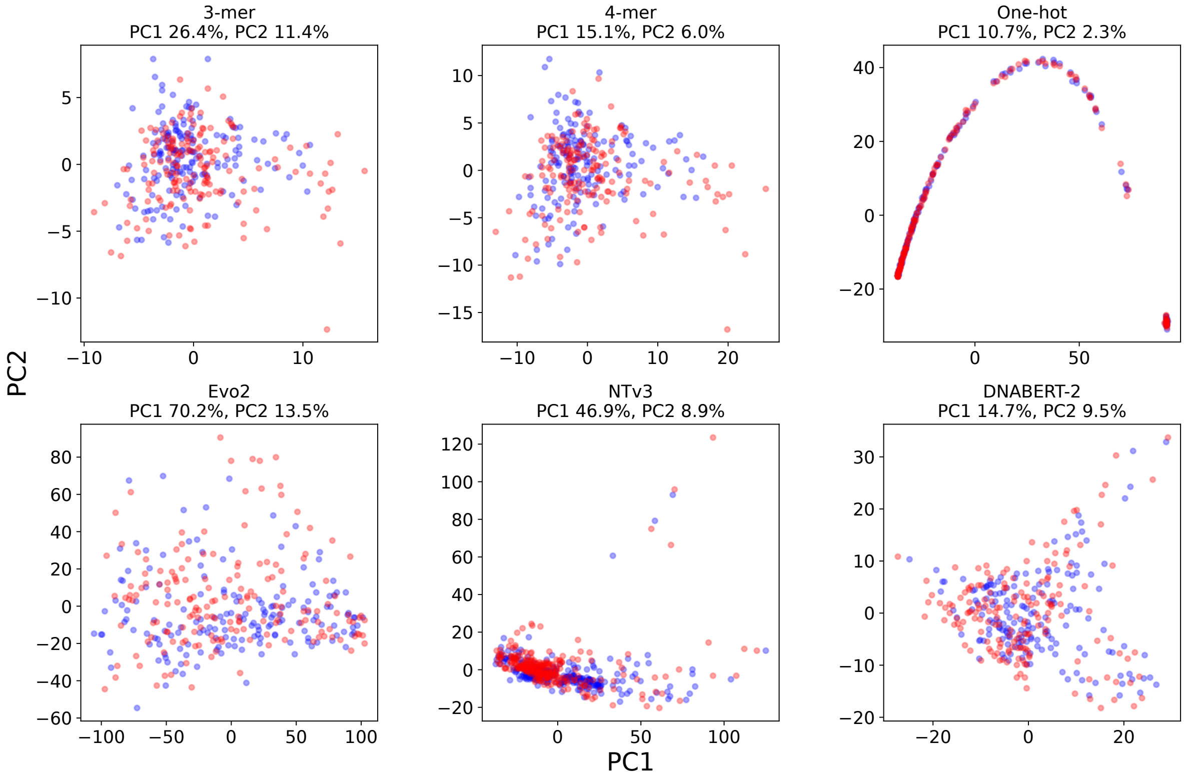


Fig. S2: **PCA visualization of sequence representations on the DRIPc-seq validation dataset.** PCA projections are shown for 3-mer, 4-mer, one-hot, Evo2, NTv3, and DNABERT-2 representations using the DRIPc-seq validation dataset from R-loop Tracker. Positive R-loop sequences are shown in red, and adjacent matched negative sequences are shown in blue. The dataset contains 170 positive and 168 negative sequences. PCA was used instead of UMAP for this smaller validation set to provide a more stable and directly interpretable low-dimensional visualization.

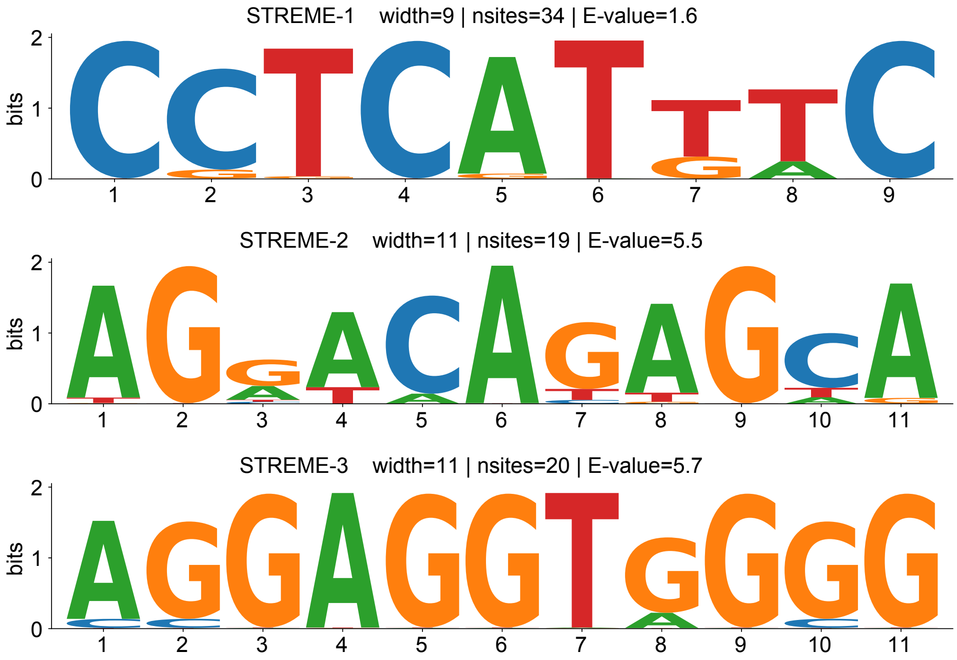


Fig. S3: **Candidate enriched sequence motifs identified from the DRIPc-seq validation dataset.** Sequence logos are shown for the top three candidate motifs identified by STREME using the DRIPc-seq validation dataset from R-loop Tracker. Positive R-loop sequences were compared against adjacent matched negative sequences. The dataset contains 170 positive and 168 negative sequences. For each motif, the motif width, number of contributing sites, and E-value are shown above the corresponding logo.
